# Population codes of prior knowledge learned through environmental regularities

**DOI:** 10.1101/688630

**Authors:** Silvan C. Quax, Sander E. Bosch, Marius V. Peelen, Marcel A. J. van Gerven

## Abstract

How the brain makes correct inferences about its environment based on noisy and ambiguous observations, is one of the fundamental questions in Neuroscience. Prior knowledge about the probability with which certain events occur in the environment plays an important role in this process. Humans are able to incorporate such prior knowledge in an efficient, Bayes optimal, way in many situations, but it remains an open question how the brain acquires and represents this prior knowledge. The long time spans over which prior knowledge is acquired make it a challenging question to investigate experimentally. In order to guide future experiments with clear empirical predictions, we used a neural network model to learn two commonly used tasks in the experimental literature (i.e. orientation classification and orientation estimation) where the prior probability of observing a certain stimulus is manipulated. We show that a population of neurons learns to correctly represent and incorporate prior knowledge, by only receiving feedback about the accuracy of their inference from trial-to-trial and without any probabilistic feedback. We identify different factors that can influence the neural responses to unexpected or expected stimuli, and find a novel mechanism that changes the activation threshold of neurons, depending on the prior probability of the encoded stimulus. In a task where estimating the exact stimulus value is important, more likely stimuli also led to denser tuning curve distributions and narrower tuning curves, allocating computational resources such that information processing is enhanced for more likely stimuli. These results can explain several different experimental findings and clarify why some contradicting observations concerning the neural responses to expected versus unexpected stimuli have been reported and pose some clear and testable predictions about the neural representation of prior knowledge that can guide future experiments.

## Introduction

During perception, the brain is continually trying to infer the cause of the perceptual input that reaches the senses. Efficient neural machinery is required to successfully perform this inference given the noise and ambiguity in our observations. Prior knowledge about the occurrence of stimuli in our environment can help to make better perceptual decisions about these noisy and ambiguous observations. The influence of prior knowledge on the perceptual decisions made by humans and animals has been demonstrated in many situations.

For example, the fact that light almost always comes from above, influences our perception of ambiguous shapes^1,2^ and our prior knowledge of shadows causes equally bright surfaces to appear of different brightness^3^. Similarly, the perception of oriented bars shows a bias towards the cardinal axis, the so called ‘oblique’ effect, which has been successfully explained by an optimal environmental prior which accounts for the frequency with which certain orientations occur in the natural environment^4^. Furthermore, owls systematically underestimate the direction of sound sources occurring far away from the center of their gaze, which has been explained as a bias induced by a prior that emphasizes directions near the center of gaze^5^.

The prior knowledge that is used in such perceptual decision making should somehow match the environmental statistics in order to enable optimal inference. How the brain learns to represent such prior knowledge and incorporate it in its sensory processing, is still an open question. While one possibility is that prior knowledge is transferred through evolutionary means, this would lead to very inflexible priors, not optimally adapted to changing environments. Alternatively, the brain learns to represent prior probabilities about the environment during its lifetime through the frequency with which an object is observed over time^3^.

Different ideas exist about the representation of prior probabilities in the brain. One popular theory is that uncertainty about a variable is represented together with the variable itself, by the population activity of Poisson neurons^6^. Alternatively, the variability and the mean of neural responses can be representing separate aspects of a posterior probability distribution, such that the uncertainty encoded by the variability of neural firing is not represented in the mean responses of the neurons^7^. This way, prior knowledge can already be represented in the neural variability without sensory input^8^. Another theory is that specialized neurons encode prior probabilities and sensory evidence^9^.

It has also been suggested that prior knowledge is represented in the specific distribution of tuning curves in the brain^10^, where neurons are more sensitive for stimuli that have a higher prior probability. Various evidence exists in favour of these different implementations and different representations might occur in different brain areas or for different types of stimuli.

To gain more insight in the kind of representations of prior knowledge that could emerge in neural populations, we trained generic neural networks to perform two commonly-used tasks in the experimental literature. These networks can provide valuable insight into possible representations in neural populations. While it has been shown that generic neural networks can learn to perform probabilistic tasks that depend only on the current input, without a need to incorporate prior knowledge^11^, it remains an open question to what extent these networks can learn to represent a statistically optimal prior and combine this with probabilistic input.

Here, we simulated a classification task and a stimulus estimation task, where the prior probability of observing a stimulus was manipulated. Both tasks use a fundamentally different prior; a discrete prior for the different classes in the classification task and a continuous prior for the stimulus estimation task. We show that a model population of neurons learns to correctly represent and incorporate prior knowledge in both, by only receiving feedback about the accuracy of their inferences and without probabilistic feedback. We find an opposing effect of prior and input uncertainty on the sparseness of neural firing. In the classification task, prior knowledge lowers the activation thresholds for the more likely class, causing neurons to fire more easily. In the continuous task, a sharper prior leads to a shift and narrowing of tuning curves, as often observed in experimental data. These results lead to testable predictions that can help validate whether prior knowledge is represented similarly in humans and animals.

## Methods

### Neural population model

Our neural population model has a feedforward architecture consisting of an input layer, a hidden layer and an output layer (Fig 1). The input layer consists of a group of 50 independent Poisson neurons that respond to a stimulus, *s*, with firing rates, **r** = [*r*_1_, …, *r*_*N*_], each generated from its own independent Poisson distribution^11^. The probability that neuron *i* responds with firing rate *r*_*i*_ is given by

**Figure 1.**
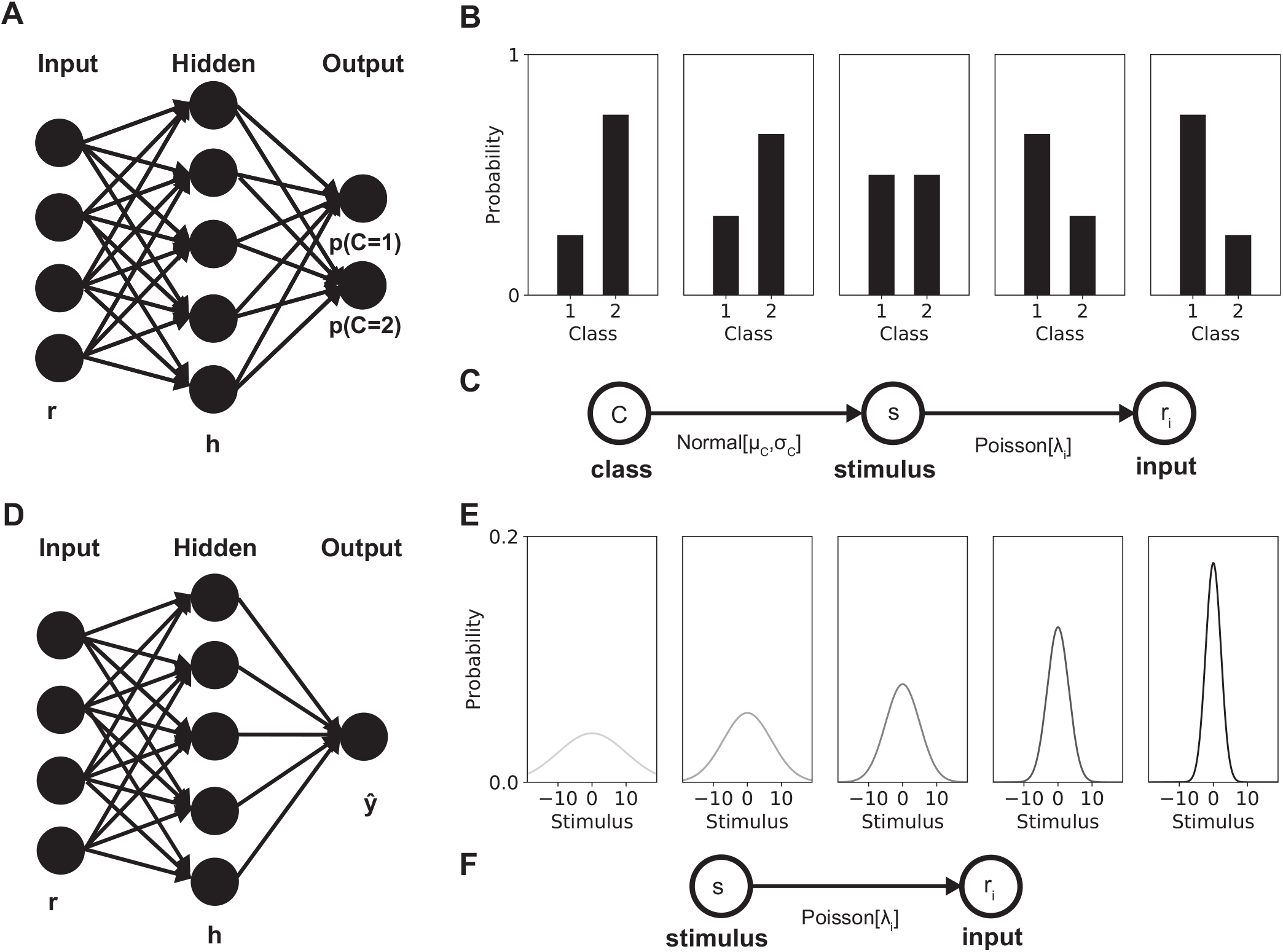
Neural population model and task overview. Two different tasks were studied, a classification task (top row) and a stimulus estimation task (bottom row). The neural network for the classification task and estimation task had a similar input and hidden layer, only the output layer differed. **(A)** Two softmax outputs (one for each class) indicated the probability that the stimulus observed came from the respective class in the classification task. **(B)** The stimuli from each class came from a normal distribution with a mean and variance specific to that class. Different prior probabilities for the classes were tested as represented by the bars. **(C)** The generative model used to generate a stimulus from a selected class and subsequently generate firing rates for the input Poisson neurons. **(D)** A single output indicated the estimated stimulus value in the estimation task. **(E)** The stimuli presented to the network were drawn from a normally distributed prior. Priors with different variances were tested: 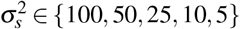 (indicated by gray scale, from left to right). **(F)** The generative model for the estimation task, used to generate firing rates for the input Poisson neurons from a generated stimulus.

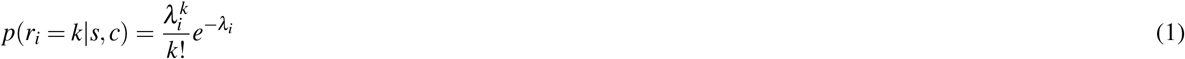

where the rate parameter *λ*_*i*_ is determined by Gaussian tuning curves *f*_*i*_(*s*), specific for each input neuron, scaled with a global contrast parameter *c*, i.e. *λ*_*i*_ = *c f*_*i*_(*s*). The Gaussian input tuning curves for each neuron are thus given by

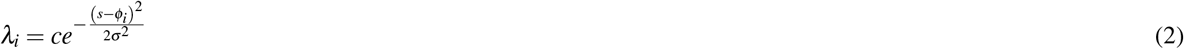

where the maxima of the tuning curves, *ϕ* = [*ϕ*_1_, …, *ϕ*_*N*_], were uniformly distributed over a range of −20 to 20 and the width of every tuning curve was given by *σ* ^2^ = 10. The contrast parameter, *c*, determined the quality of the stimuli. The range of contrasts used depended on the task and is specified in the task descriptions below.

The hidden layer was densely connected to the input layer with weights, **W**, and consisted of, *K* = 200, rectified linear units (ReLUs) with bias parameters *b*^*k*^ and firing rates **h** = [*h*^1^, …, *h*^*K*^]

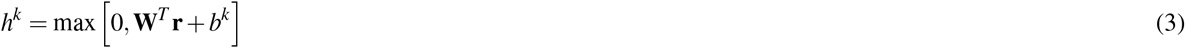

for 1 ≤*k* ≤K.

The output layer of the network was densely connected to the hidden layer via output weights, **U**, and consisted either of two units, one for every class, in the classification task (Fig 1A), or a single unit, in the stimulus estimation task (Fig 1D). A softmax function was applied to the output units of the classification task to convert them to probabilities.

In some experiments a cue was added to indicate which class was more likely to occur on a certain trial. Two important ways in which a cue can modulate a neural population to represent prior knowledge are through gain modulations or biases. To enable the network to potentially learn such representations, we equipped every ReLU unit with both a gain parameter, 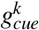 and an extra bias parameter, 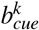, per unit, that could be modulated by the cue. These extra parameters were also learned via the backpropagation algorithm. That is,

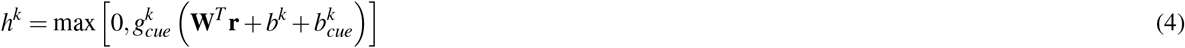

for 1 ≤*k* ≤K.

### Network training

All simulations were performed in Python. To train our neural networks we used the PyTorch neural network library^12^. Parameters were initialized from a uniform distribution in 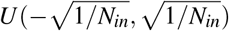, with *N*_*in*_ the number of inputs into a layer. Thus for **W** and **b**, *N*_*in*_ = 50 and for **U**, *N*_*in*_ = 200. In the experiments where a cue was added both **g**_*cue*_ and **b**_*cue*_ were initialized the same way with *N*_*in*_ = 50.

The networks were trained using the backpropagation algorithm. To optimize the network parameters, **W, b, U** and in case of the cue experiments **g**_**cue**_, **b**_**cue**_, we used Adam^13^, with a learning rate of 2 *×* 10^−4^. We used a minibatch size of 10 per iteration. Every network was trained for 100 epochs with 1000 iterations per epoch. Stimuli were presented in a random order during training.

### Classification task

To study the representations of prior knowledge emerging in a population of neurons, we designed a classification task. Stimuli could be drawn either from class 1 or class 2. The relative frequency with which a stimulus was drawn from a class determined the prior probability with which stimuli from this class occurred. The stimulus was converted to firing rates by a population of Poisson neurons, which in turn served as the input to a population of hidden rate-based neurons, which was read out by a linear layer.

In this classification task, the network had to decide whether observed stimuli, *s*, belonged either to the first class (*C* = 1) or the second class (*C* = 2). Both classes were determined by a one-dimensional Gaussian distribution with the same variance, 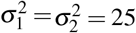, but different means, with *μ*_1_ = −5 and *μ*_2_ = 5 (Fig 1B,C). Stimuli with contrast values *c* ∈ {0.5, 1.2, 1.9, 2.6, 3.3, 0} were presented. We used five different priors over the two classes chosen from *π* ∈ {0.25, 0.33, 0.5, 0.67, 0.75} such that *p*(*C* = 1) = 1 − *p*(*C* = 2) = *π*. Because of the symmetry of the binary classification problem these five different priors made up three independent ratios. However, in order to ensure that no intrinsic bias exists in our network all five options are investigated in the results section. The prior probabilities for the two classes determined the frequency with which samples from these classes were shown during training.

In the network solving the classification task, the output of the read-out layer was converted by a softmax function to obtain a normalized probability. For class *i* the probability is given by

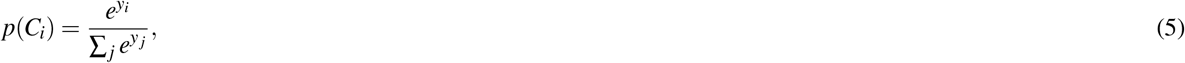

with *y*_*i*_ the read-out for class *i* and the sum representing the sum over all classes (two in this case). The loss per trial is then calculated using this softmax read-out and the binary class label of the stimulus with the binary cross entropy

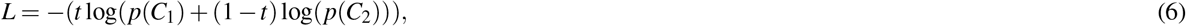

with the target label, *t*, being 1 when the class is 1 and being 0 when the class is 2. To observe how optimal our network performed this task we compared the posterior learned by the network (the softmax read-out) with the optimal Bayesian posterior probability. We determined these posterior probabilities for both classes. We wish to estimate the posterior probability of a class *C* given observations of neuronal responses **r**:

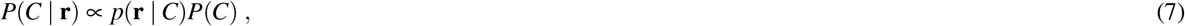

where the likelihood is given by:

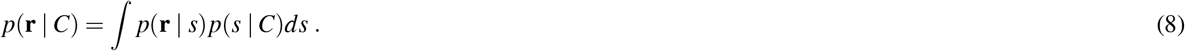

In our task we draw the stimulus from a normal distribution depending on the class:

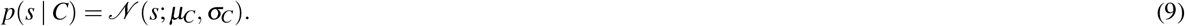

The conditional probability of neuronal responses **r** of the Poisson neurons can be written in the following form:

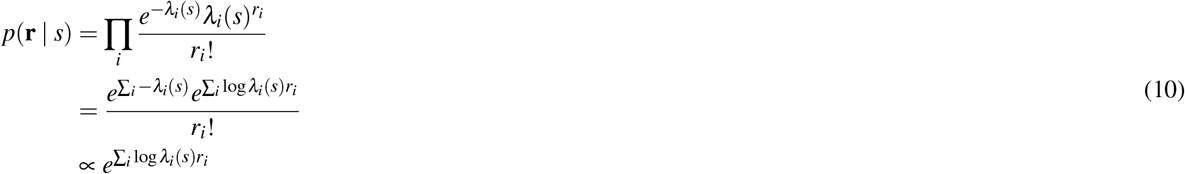

where we dropped the parts that are independent of *s* and assumed that ∑_*i*_ − *λ*_*i*_(*s*) is constant and thus also independent of *s*.

If we let the tuning functions take a Gaussian form this results in

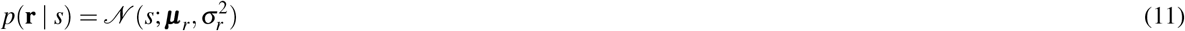

with ***μ***_*r*_ = *ϕ*^*T*^(**1**^*T*^**r**)^−1^ and 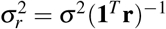

Since both parts of the likelihood are normally distributed we can marginalize the likelihood to arrive at the following form

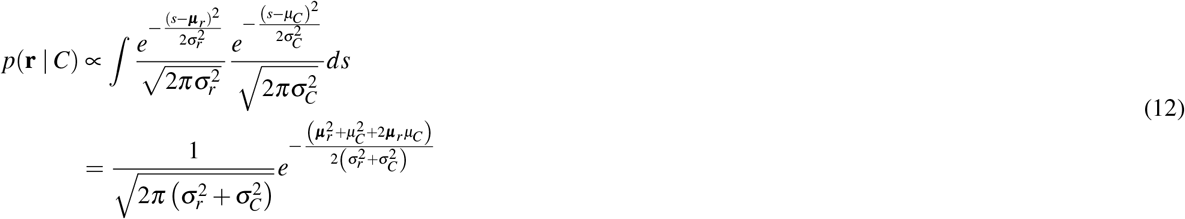

The log posterior ratio of the probability that an observation is of one of both classes is given by

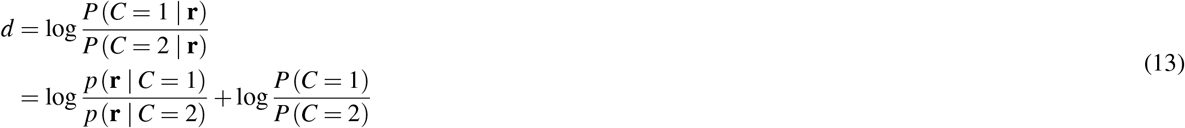

For our task substituting in Equation 12 in Equation 13 drops out the normalization term and gives

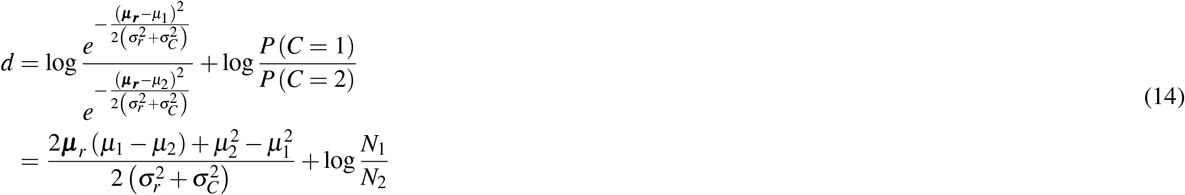

where *μ*_*c*_ is the mean response of class *c* and *N*_*c*_ is the number of training samples that have been shown for class *c*.

The posterior probability for class 1 can then be determined by

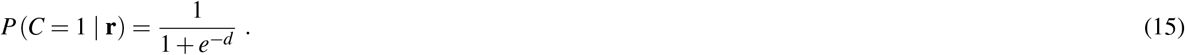

### Estimation task

In the estimation task, the network had to estimate the continuous value of the observed stimulus *s*. Stimuli were drawn from a Gaussian prior distribution with mean *μ*_*s*_ = 0 and variance 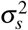 (Fig 1E,F)

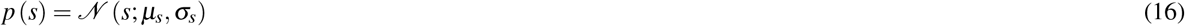

Priors with different variances were tested: 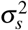∈ {100, 50, 25, 10, 5}. Stimuli with contrast values *c* ∈ {0.30, 0.72, 1.45, 2.26, 2.86, 3.2} were presented.

The network solving this task had the same basic architecture as for the classification task. Only the readout was in this case not converted by a softmax function, but directly reflected an estimate of the stimulus value. The mean squared error loss between the estimated stimulus value, *ŝ*, and actual stimulus value was used to optimize the network for this task

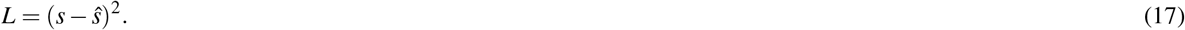

To determine the optimal Bayesian estimate, we would like to estimate the most probable stimulus value *s*, given observations of neuronal responses **r**

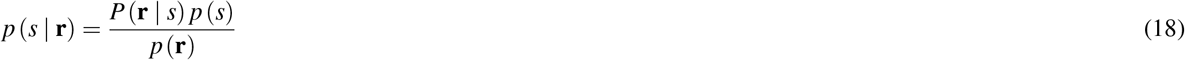

The response to a certain stimulus value is again given by Poisson neurons (Eq 10-11). The posterior will then be proportional to the product of two normal distributions, which will again be normally distributed

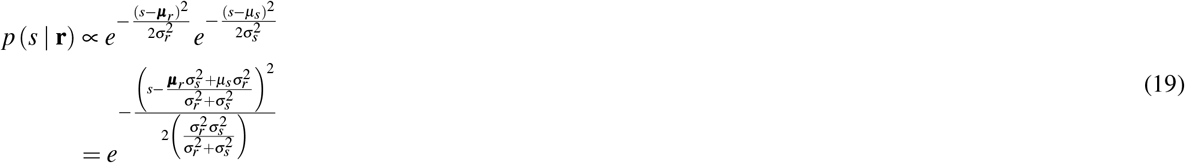

The posterior mean is given by the maximum-a-posteriori (MAP) estimate, which is used as optimal Bayesian estimate of the underlying stimulus value and compared against the stimulus estimate given by the network

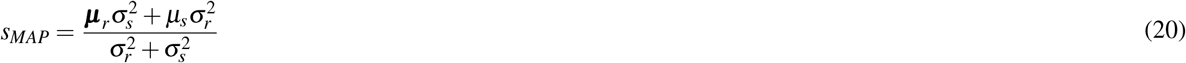

## Results

To study the representations of prior knowledge emerging in a population of neurons, we designed two representative tasks with a distinctive type of prior covering many behavioral experiments. In the first task the network had to decide to which of two classes a stimulus belonged. A discrete prior determined the prior probability of observing a stimulus from one of these two classes. In the second task the network had to estimate the value of a stimulus. Here a continuous prior determined the prior probability of observing a certain stimulus value (see Methods section for further details on the different tasks). Since the prior affects the optimal outputs in both tasks differently, there could be distinctive representations emerging, making both example tasks worth investigating. In the first part of the results we investigate the classification task, while the latter part of the results focuses on the estimation task.

### Optimal prior learned on trial-by-trial basis

To investigate whether the network was able to learn a prior about the occurrence of the different classes and incorporate this optimally, we compared a linear softmax readout of the network with an optimal Bayesian solution of the task (Fig 2).

**Figure 2.**
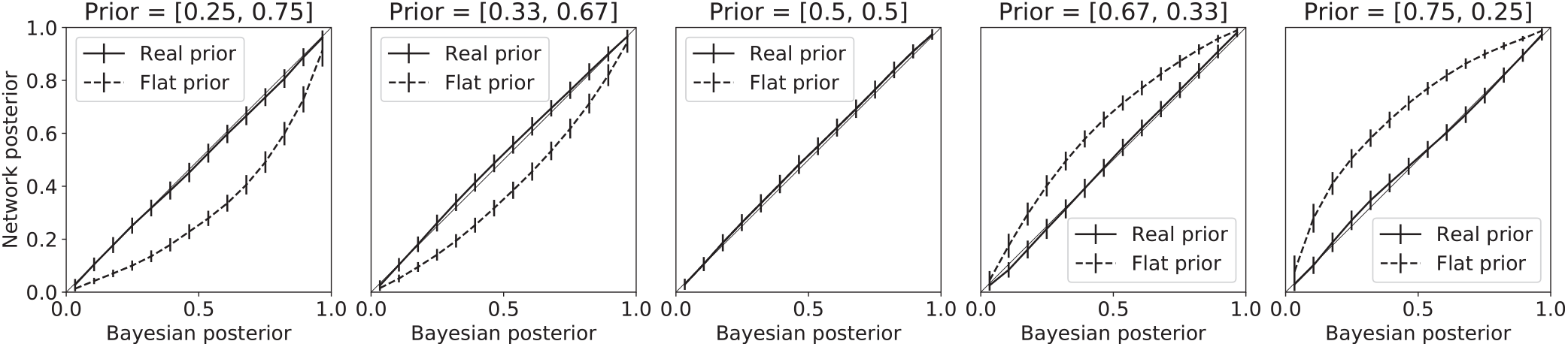
Network posterior matches optimal Bayesian estimates. The neural network learned to choose each class with a certain probability by getting feedback about correct or incorrect classification. Based on Bayesian statistics the optimal posterior estimate depends on the prior probability of a class occurring. This prior probability was influenced by the relative frequency that a stimulus from a class was shown during training. Five prior probability ratios were tested: [0.25, 0.75], [0.33, 0.67], [0.5, 0.5], [0.67, 0.33] and [0.75, 0.25] for class 1 and class 2 respectively. The posterior that the network learned is plotted against the optimal Bayesian estimates that accounts for the prior (solid line), or that ignores the prior probability, i.e. a uniform prior (dashed line). The network posterior matches the optimal Bayesian estimate that accounts for the prior and thus learns to correctly incorporate prior knowledge in its connections. Error bars represent standard deviation over the validation trials.

**Figure 3.**
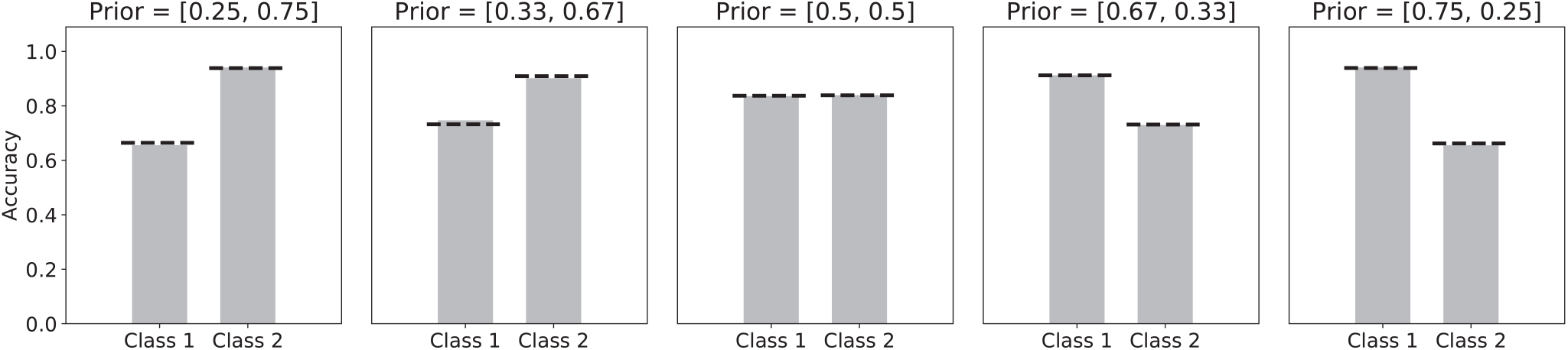
Prior enhances accuracy for more likely class. The accuracy with which stimuli were correctly classified was influenced by the prior probability of a certain class. A uniform prior (panel 3) led to equal accuracy on stimuli from both classes. A network trained on more stimuli from class 2 than class 1 (panel 1 and 2) showed higher accuracy on class 2 stimuli, but lower accuracy on class 1 stimuli, compared to the uniform prior. On the other hand a network trained on more stimuli from class 1 than class 2 (panel 4 and 5) showed higher accuracy on class 1 stimuli, but lower accuracy on class 2 stimuli, compared to the uniform prior. (Dashed line represents accuracy of optimal Bayesian model.

While the network did not receive any probabilistic feedback about the prior probabilities of either class, it formed a correct hidden representation of these prior probabilities and adjusted its choices in a Bayes-optimal way. If the network would only perform a maximum likelihood estimation (i.e. use a uniform prior), the network posterior would have matched the dashed line (see Fig 1). This result suggests that no higher order cognitive functions that keep track of the relative frequency by which an object class is observed are needed to form a Bayes optimal prior in the brain.

The accuracy of the classifications (as quantified by the fraction of correctly classified labels) was calculated by comparing the class label with the highest probability according to the read-out softmax of the network with the real class label. The use of this prior knowledge led to more accurate responses the class with higher prior probability, a behaviour that humans have also shown in many studies^4,14^.

### Representations of prior probability

To investigate how the neural population encoded the prior probability of the different classes, we first looked at the neural activation in response to different stimuli. Both uncertainty in the input neurons and in the prior probability of the different classes were manipulated. Less uncertainty in the input led to sparser population responses, as previously reported by^11^) (Fig 4A). However, when the posterior uncertainty is considered, we observed that more certainty about the presented class actually led to stronger responses, where more neurons fire. Uncertainty in the input strongly changes responses to stimuli that lead to uncertain posterior responses (around 0.5).

**Figure 4.**
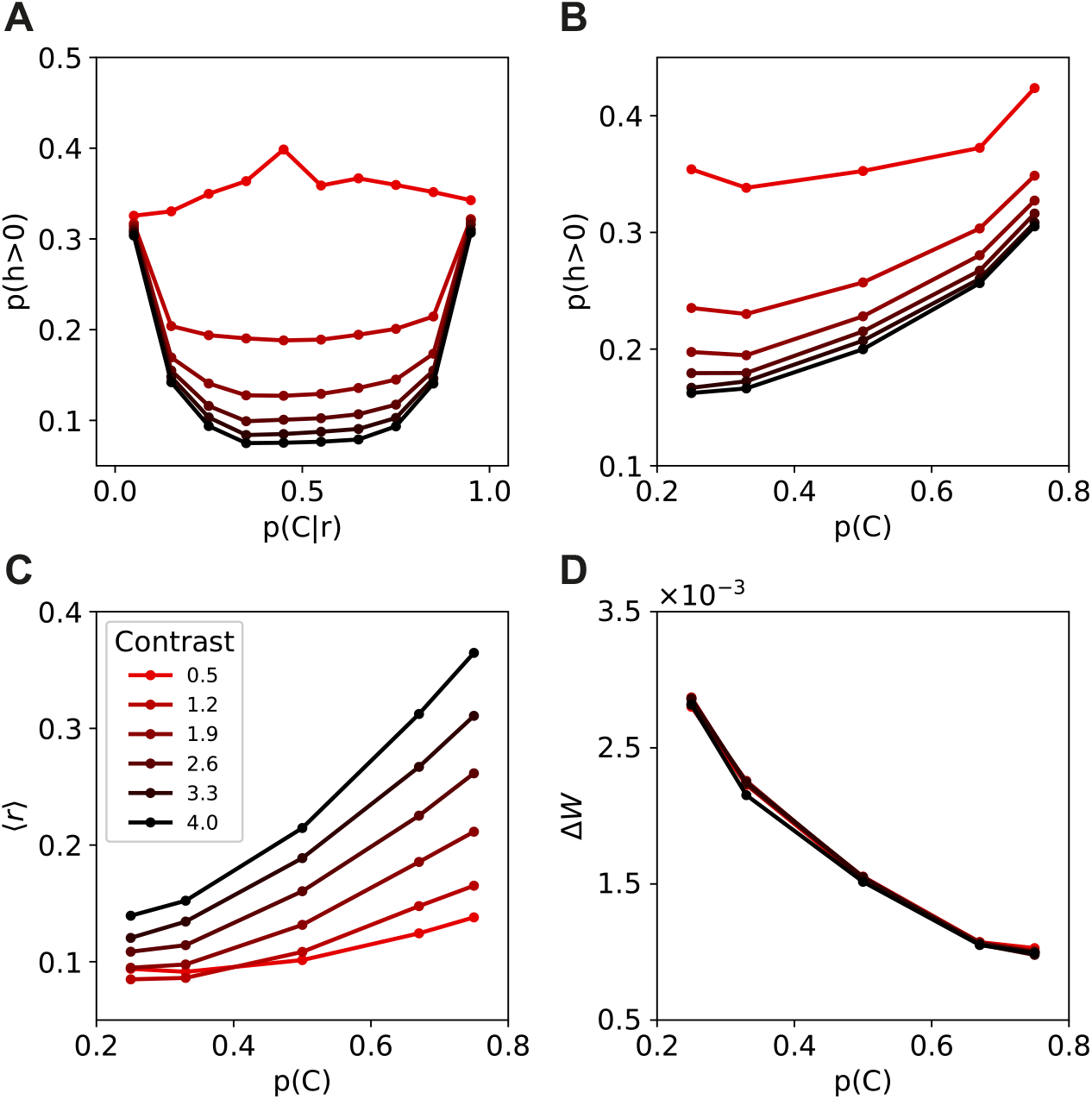
Neural activity changes due to input and prior uncertainty. **(A)** Number of neurons with non-zero activity in response to both stimuli from class 1 and class 2. Fewer neurons in the hidden layer responded when the input contrast was higher. On the other hand, more neurons responded when the posterior was certain which class was presented. **(B)** Number of neurons with non-zero activity in response to stimuli from class 1. While fewer neurons responded to higher contrast stimuli, more neurons were active when the prior probability of the presented stimulus class was higher. **(C)** The average activity of the population increased both with stimulus contrast and prior probability. **(D)** The weights in the network require less adaptation in response to likely stimuli, while the contrast has no influence on the strength of the weight updates.

These responses could be explained by a template matching code, where the neural population develops templates for both classes against which the input is matched. Very certain inputs led to clear matches with either of the classes, or a clear mismatch when a stimulus came from the decision boundary. On the other hand, more noisy inputs led to partial matches with both templates, even for stimuli around the decision boundary.

Next, we considered the effect of uncertainty in the prior probability of the classes. A higher prior probability for a certain class led to more neurons firing in response to stimuli of this class. Uncertainty in the input and uncertainty in the prior thus have opposing effects on the sparseness of the neural responses. While a higher contrast in the input (less uncertainty) led to sparser neural responses, a higher prior probability for a class (also less uncertainty) led to less sparse neural responses for this class (Fig 4B). The average firing rate for the entire neural population increased both with more certainty in the input, as well as with more certainty in the prior (Fig 4C). On the other hand, decreased the learning signal in response to more likely stimuli, requiring less adaptation of the weights (Fig 4D).

### Prior probability encoded by changing activation thresholds

The neural responses of the population offer measurable predictions that can be experimentally validated. A more difficult, but nonetheless interesting, question to answer experimentally is which mechanism gives rise to these neural responses. Changes in the properties of individual neurons and wiring between neurons in the population should somehow represent the prior probability of diverse stimuli.

To see how the parameters in our network encoded this prior information we determined for every neuron in the population whether it contributed more to class 1 or class 2. We analyzed how the weights of input to hidden units, **W**, the bias of the hidden units, **b**, and the weights from hidden to output units, **U**, changed with different prior probabilities for a class. The neurons in the population responsive to the class with higher prior probability developed on average a higher bias, effectively lowering their activation threshold (Fig 5A). Lower thresholds explain the fact that more neurons fire when a class has higher prior probability.

**Figure 5.**
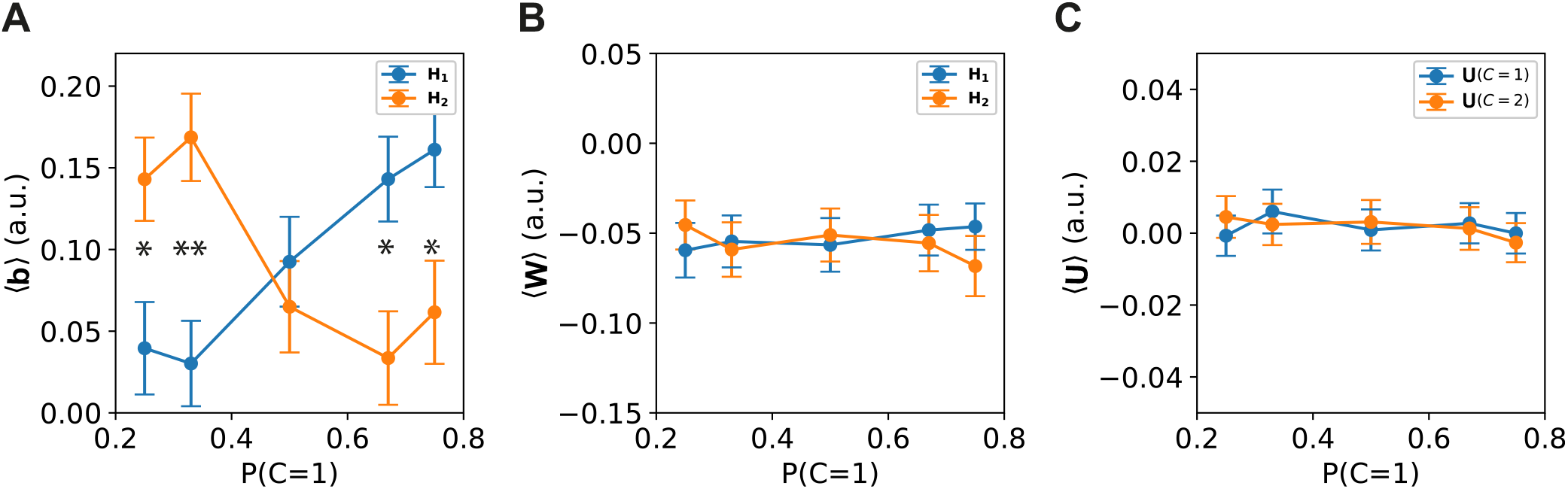
Prior probability encoded by changing activation thresholds. The hidden units could change the decisions of the output units using the bias parameters (**b**), hidden weights (**W**) or output weights (**U**). For every hidden neuron, it was determined whether it contributed stronger to class 1 or class 2 by comparing the output weights to class 1, **U**(*C* = 1), against the output weights to class 2, **U**(*C* = 2). That resulted in two subpopulations of hidden units: **H**_1_ = **H**[**U**(*C* = 1) *>* **U**(*C* = 2)] and **H**_2_ = **H**[**U**(*C* = 1) *<* **U**(*C* = 2)]. The biases and weights of the hidden units were averaged accordingly and are plotted for different prior probability ratios (*P*(*C* = 1) ∈ [0.25, 0.33, 0.5, 0.67, 0.75]). **(A)** Only the biases of the hidden neurons changed as a result of different prior probabilities, where a higher probability of a class led to higher bias values in neurons coding for this class, thus lowering the activation threshold in these neurons. **(B)** The hidden weights did not change with prior probability. **(C)** Neither did the output weights. Error bars represent standard error over the units. (* *p <* 0.01, ** *p <* 0.001).

As a control we also checked for the readout weights of the neural population, but found no effect of the prior probability on the mean of the readout weights to either class (Fig 5C).

### Prior probabilities indicated by contextual cues

In our everyday environment the prior probability of a stimulus occurring often depends strongly on the context. Since in these situations the context has to be able to manipulate the prior on a very short time scale, it is not possible that long term synaptic changes underlie such representations. Different hypotheses exist about the neural mechanisms underlying the encoding of prior knowledge dynamically in such cases. A popular hypothesis is that prior knowledge is encoded by modulating the gain of neurons. Neurons encoding for a more probable stimulus fire more strongly in response to this stimulus. Another possibility is that prior knowledge modulates the activation threshold of neurons. A neuron encoding a more probable stimulus is then more easily activated in response to this stimulus. In biological networks, a lower activation threshold can be achieved by excitatory modulation, while an higher activation threshold is achieved by inhibitory modulation, either of the shunting or hyperpolarizing type^15,16^. Gain modulation on the other hand requires an additional background noise input that is a combination of coupled excitatory and inhibitory input^16^. Both these mechanisms have been widely observed in the brain^17,18^ and could play an important role in representing prior knowledge.

To see if our network is able to learn such a prior that has to change flexibly depending on the context, we added an extra cue input feeding into the hidden layer, with connections to both the gain and bias of the neurons. A single neural network was trained that had to learn to change its prior probabilities depending on the cue. Given the cue the probability of observing a stimulus from class 1 was either 25% or 75% (vice versa for class 2). The network was able to learn to correctly account for the prior probabilities for the different context cues (Fig 6A,B). Since all network parameters are the same for both cue conditions, the network had to learn a flexible mechanism to account for the prior probability. Closer inspection of the effect of the cue on the hidden neuron input shows that the hidden neurons encoding the more likely class received both a stronger baseline input from the cue (Fig 6C) and had a stronger gain (Fig 6D).

**Figure 6.**
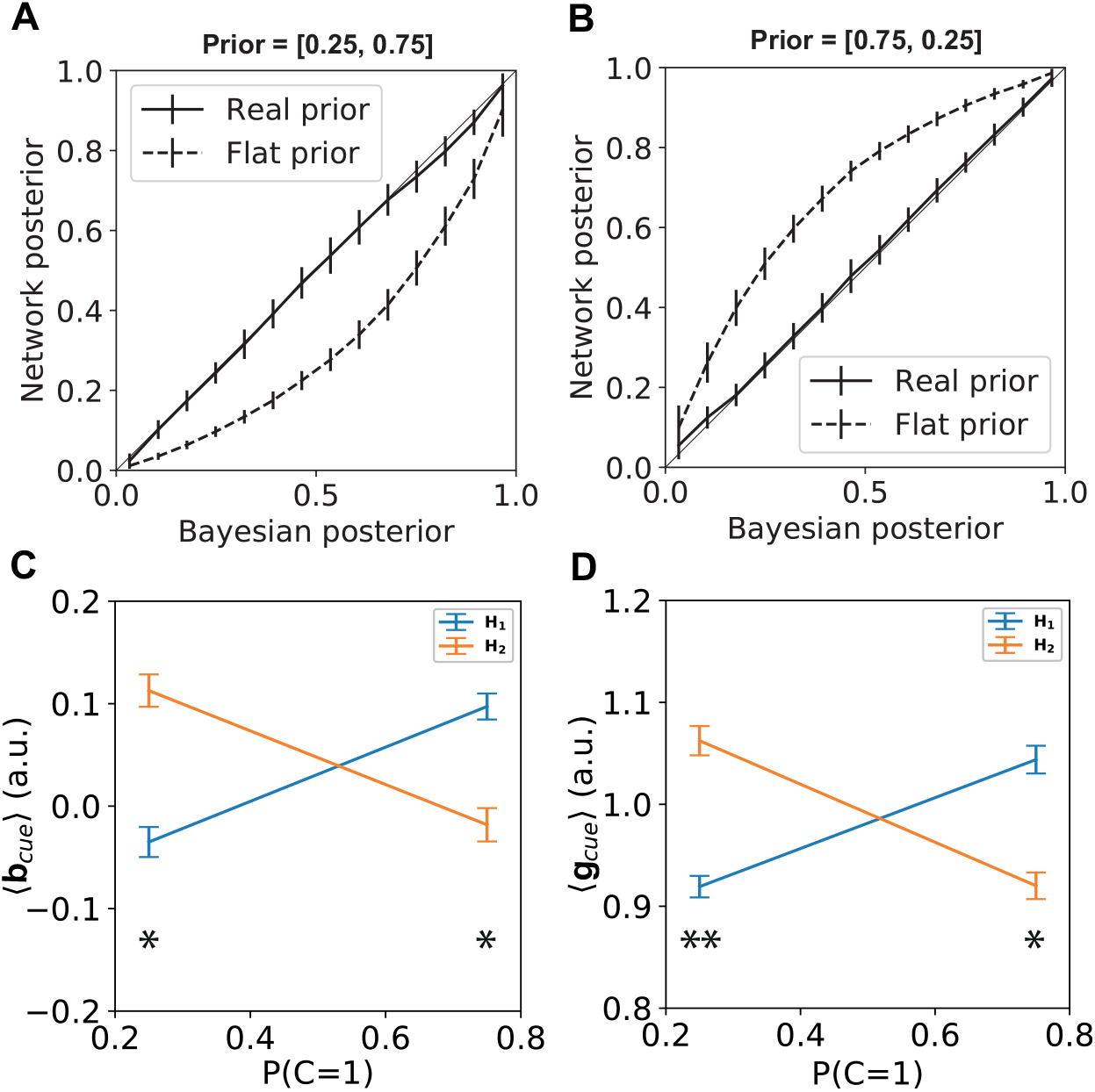
Contextual cue modifies activation thresholds. The frequency with which each class of stimuli was shown depended on a contextual cue. Depending on the cue the prior probability of class 1 and class 2 could either be [0.25, 0.75] or [0.75, 0.25]. **(A**,**B)** The network learned to adapt its output probability to reflect the optimal Bayesian estimate given the cue. Error bars represent standard deviation over the validation trials. **(C**,**D)** For every hidden neuron, it was determined whether it contributed stronger to class 1 or class 2. The input from the cue to the bias and gain of the hidden neurons was averaged accordingly for both both cues. The cue for the context where class 2 is more likely caused both a higher input to the bias and gain of hidden neurons encoding class 2 and a lower input to the hidden neurons encoding class 1. The opposite effect occurred for the cue for the context where class 1 is more likely. Error bars represent standard error over the units. (* *p <* 10^−7^, ** *p <* 10^−10^).

Since both the bias and the gain are influenced by the cue, we investigated whether one of these played a more important role in representing the correct prior knowledge. We analyzed the accuracy of the network on classifying both classes under the different cues. When the cue influenced both the gain and the bias, the prior caused a large difference in accuracy between both classes (Fig 7, blue bars). Next we turned off either the influence of the cue on the gain (Fig 7, orange bars) or the influence of the cue on the bias 7, green bars). Turning off the influence of the cue on the gain diminished the effect of the prior on the accuracy only slightly. Turning of the influence of the cue on the bias on the other hand, diminished the effect of the prior on the accuracy almost completely. The bias thus seems to play a greater role in representing the prior knowledge.

**Figure 7.**
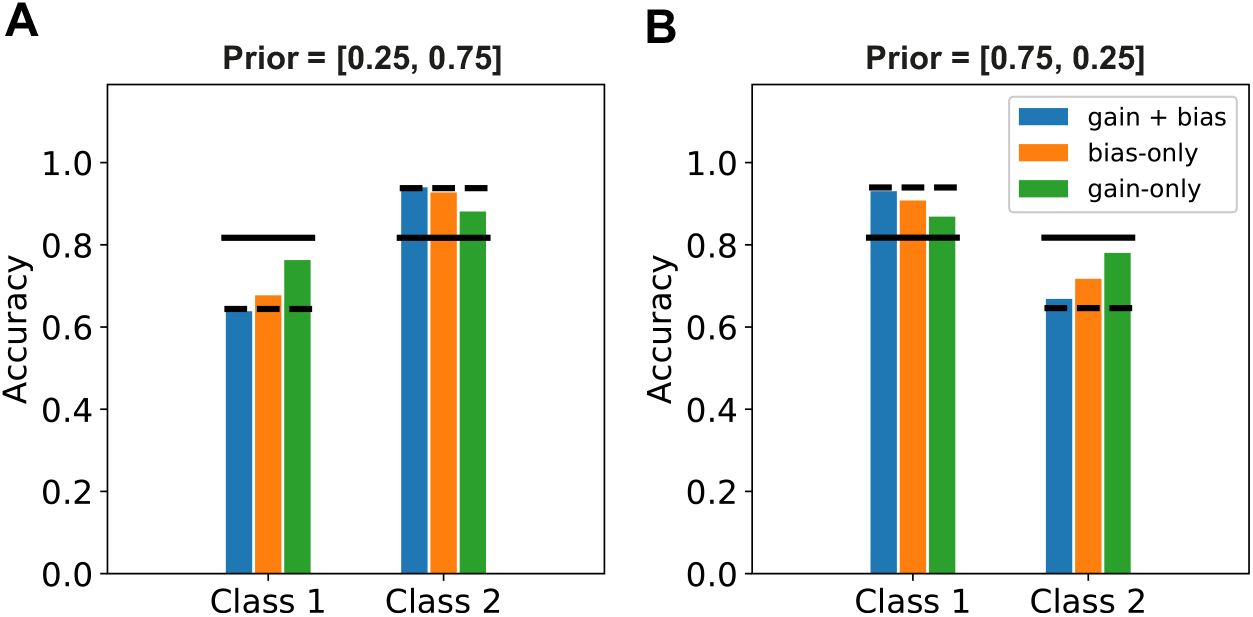
Cue influence mainly through bias. The accuracy for both classes was determined for the cued priors (left, [0.25, 0.75], right, [0.75,0.25]). **(A**,**B)** The class with the highest prior probability according to the cue, was decoded much more accurately when the cue influenced both the gain and the bias (blue bars), closely resembling the accuracy of the optimal Bayesian model (dashed line). Turning off the cue effect on the gain diminished this effect only slightly (orange bars). Turning off the cue effect on the bias diminished this effect much more, almost completely removing any effect of the cued prior (green bars), making the accuracy more similar to the accuracy of a Bayesian model that only accounts for the likelihood (solid line).

### Prior probability encoded by efficient allocation of neural resources

In the previous section we studied how prior knowledge about a discrete class can be learned and encoded in a neural population. However, many tasks in the brain require a prior representation of a continuous environmental variable. E.g., an often used example in the experimental literature is the estimation of the orientation of gratings^4^, but also sound localization has been explained this way^5^. Since the effect of such a continuous prior on the Bayesian posterior is mathematically more complex, this might also require a different neural representation. To study whether such a continuous prior can also be successfully learned on a trial-by-trial basis and whether the representations differ from the discrete case, we trained our neural population to estimate the value of a noisy stimulus. Different Gaussian priors, *𝒩* (*μ* = 0, *σ* ^2^ ∈ {100, 50, 25, 10, 5 }), were tested, where the probability that a certain stimulus value was presented to the network was determined by this prior distribution. The same neural population model with Poisson input neurons and ReLU hidden units as for the discrete case was used. Instead of a softmax output layer, we used a linear readout for the continuous case. The mean squared error loss was used to optimize the parameters of the network through backpropagation.

First we compare the performance of the model again with the Bayesian optimal posterior, to test whether it shows Bayesian optimal behaviour. The neural population learned to estimate the stimulus in a Bayesian way, correctly incorporating the prior knowledge about the relative frequency with which a stimulus was presented (Fig 8). A uniform prior or maximum likelihood estimate did not match the network output for priors differentiating strongly from uniform (Fig 8E).

**Figure 8.**
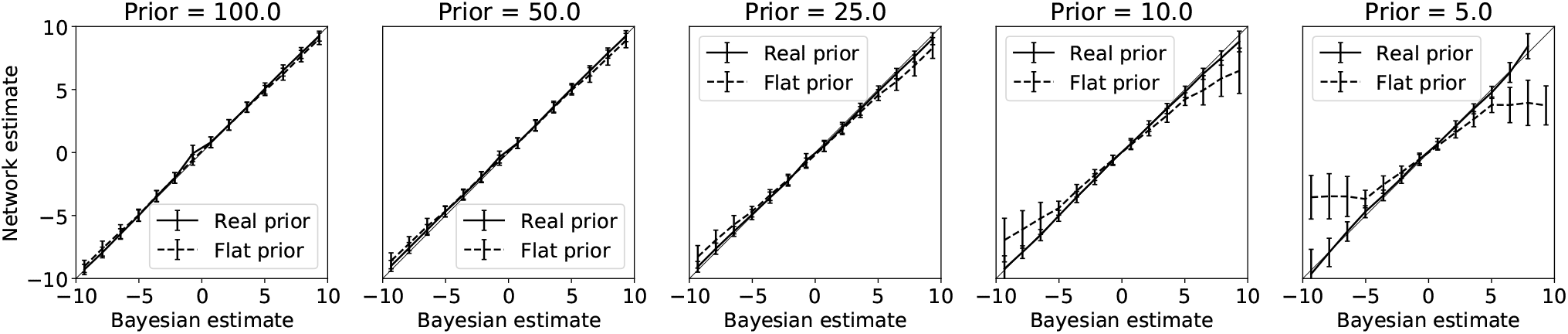
Network learns to incorporate prior knowledge optimally. Stimuli were drawn from Gaussian priors with different variances (i.e. 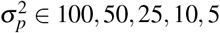 ∈ 100, 50, 25, 10, 5). During testing, stimuli in the range [−10, 10] were presented, making the 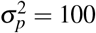 prior effectively uniform. The posterior that the network learned is plotted against the optimal Bayesian estimates that accounts for the prior (solid line), or that ignores the prior probability, i.e. a uniform prior (dashed line). While for 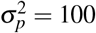 both Bayesian estimates are similar in the range [−10, 10], there is a big difference between both estimates for 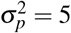. The network estimates match the Bayesian estimates accounting for the prior distribution, but not Bayesian estimates for the prior distribution. The network thus learns to incorporate the prior knowledge correctly. Error bars represent standard deviation over the validation trials.

The network showed a ‘perceptual’ bias similarly to those that are observed in humans and animals (Fig 9)^5^. The size of this bias depended on the sharpness of the prior distribution, with a sharper prior having a more pronounced bias. The bias in the network estimates matched those predicted by the optimal Bayesian estimate.

**Figure 9.**
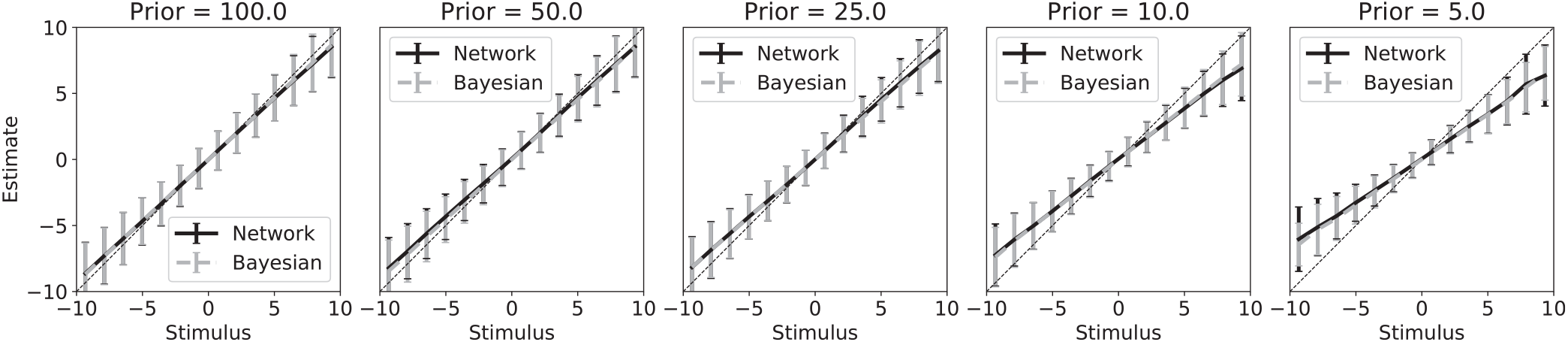
Learned prior causes perceptual bias that is statistically optimal. The network biased its estimates of the stimulus towards the mean of the prior (solid line). This effect was strongest for the sharpest prior 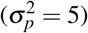. This perceptual bias matched the bias predicted by an optimal Bayesian observer (dashed line, note overlap with solid line). Error bars represent standard deviation over the validation trials.

One of the effects that is often observed in humans and animals is that the limited neural resources that are available to the organism, are used efficiently, such that more resources are allocated to ranges of stimuli that occur more often^4,5^. This improves performance for stimuli in these ranges, while consequently decreasing performance for stimuli that do not occur often. Our network showed a similar performance pattern, where stimuli around the maximum prior probability were most accurately estimated, while those further away from the maximum had a larger estimation error (Fig S1).

The neural activation patterns show a different influence of the prior for the estimation task. While a higher prior probability caused an increase in the number of neurons that fire and the overall activity in the neural population (Fig 4), we see the opposite effect in the estimation task (Fig 10A,B). Responses become sparser when the prior distribution is more narrow (Fig 10A). The effect of the stimulus contrast remains the same as for the classification task, with sparser responses on stimuli with higher contrast. The average activity in the neural population decreases when the prior distribution is more narrow (Fig 10B). The effect of the stimulus contrast on the average activity also remains the same, with higher activity in the neural population for higher stimulus contrast. For the lowest contrast value the average activity increases slightly again.

**Figure 10.**
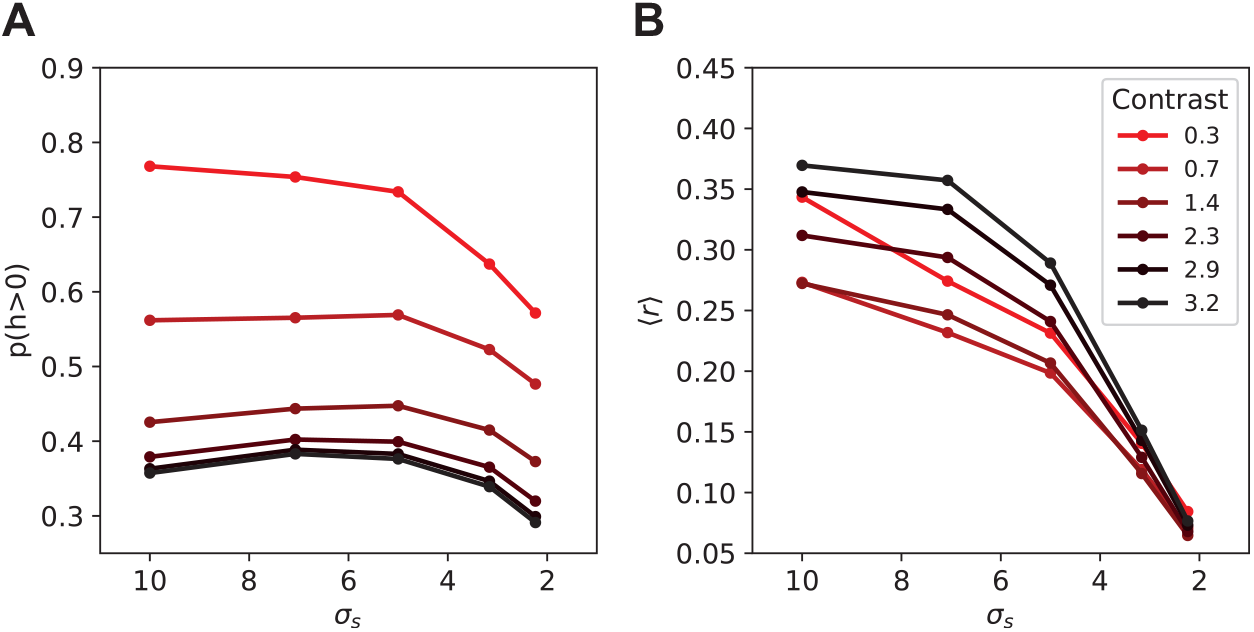
Neural activity changes due to prior uncertainty. **(A)** Number of neurons with non-zero activity in response to stimuli from different prior distributions. Fewer neurons fired when the prior distribution was more narrow. **(B)** The average activity of the population decreased with a more narrow prior probability distribution. While higher contrast led to increased neural activity. For the lowest contrast value the activity in the network also rises again.

To understand what the underlying representations are that encode this prior, we investigated the tuning curve properties of the hidden neurons. These tuning curves could change their properties such that more likely stimuli get allocated more of the limited resources. One popular hypothesis based on Gaussian tuning curves is that the density of the peaks of the tuning curves correlates with the prior probability of a stimulus, such that the highest density of tuning curves is at the stimulus value with the highest prior probability^4^. We plotted a histogram of the tuning curve peaks to see whether the highest density is found around the maximum of the prior distribution. We found a bimodal distribution where the highest density of tuning curve peaks is found slightly away from the maximum prior probability (Fig 11).

**Figure 11.**
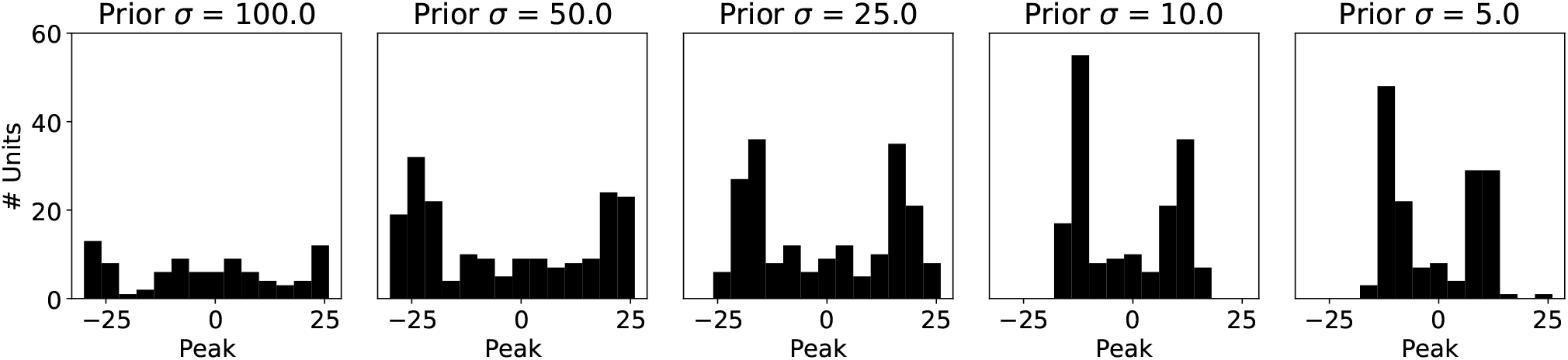
Density of tuning curves does not follow prior probability. A histogram of the peaks of all hidden neuron tuning curves is shown for the different prior conditions. While the distribution of tuning curve peaks becomes denser for the sharper prior (right panel), there is not a direct match with the Gaussian prior probability density. The tuning curve peaks rather concentrate around the flanks of the prior probability distribution.

One possible explanation is that the neural population learns to allocate the most sensitive parts of the tuning curves to the most probable stimuli range. This is part of the tuning curve where the slope is steepest, leading to the highest Fisher information. To test this alternative, we determined the slopes of the tuning curves by taking the second order derivative of the tuning curves and calculating the maximum of this slope. The histogram of these maximal slope distributions closely matches the underlying prior probability, indicating that the neural population indeed optimizes its tuning curves according to this criterion (Fig 12).

**Figure 12.**
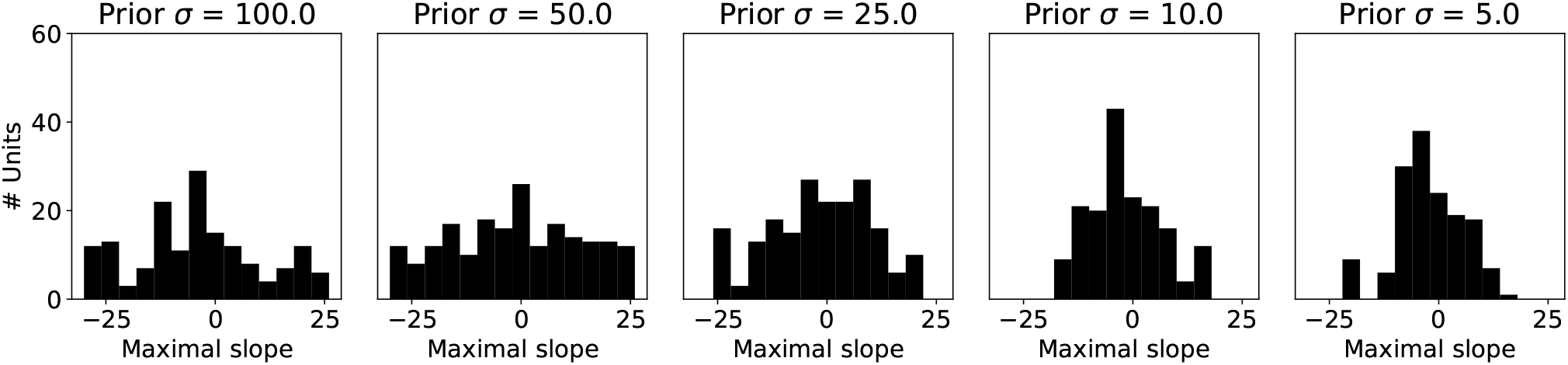
The slope of the tuning curves follow the prior distribution. A histogram of the location of the maximal slope (region most sensitive to stimulus changes) of every tuning curve is plotted for the different prior distributions. The density of maximal slopes concentrates more for the sharper prior probability distribution (right panel), forming a Gaussian bell shape.

Besides shifting tuning curves to position their maximal slope around more likely stimuli values, the tuning curves themselves can be made sharper in order to get more accurate responses around more likely stimuli values. We measured the width of the tuning curves as the full-width-at-half-maximum. The distribution of tuning curve widths shows a clear narrowing of the tuning curves for the prior distribution with the lowest variance (Fig 13), which has also been reported by theoretical derivations of the optimal tuning curve distributions for a certain prior distribution over sensory variables^10,19^. Narrower tuning curves for more likely stimuli can result in the sparser activations observed in Fig 10A.

**Figure 13.**
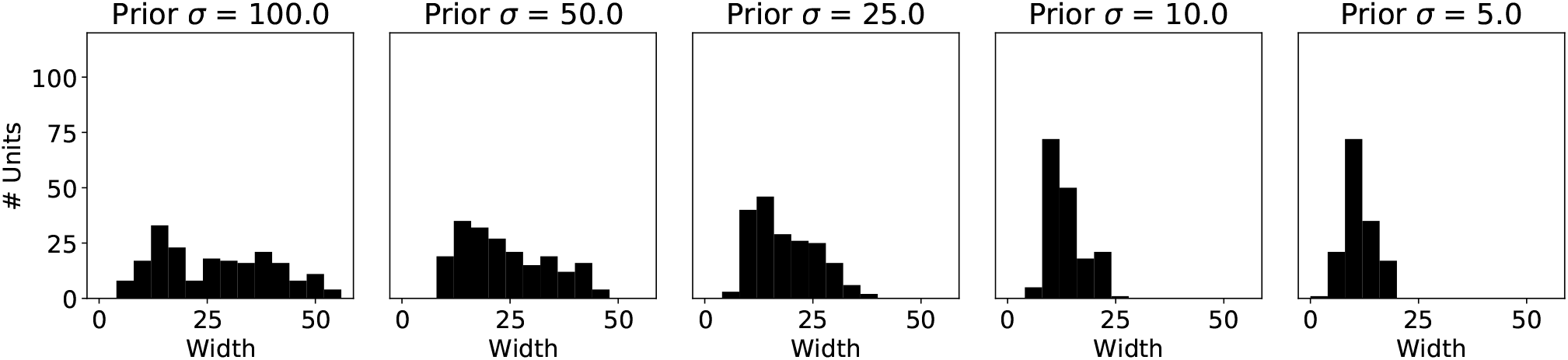
Tuning curves become narrower for sharper prior. A histogram of the full-width-at-half-maximum for all tuning curves is shown. For a sharper prior (right panel) tuning curves became narrower, effectively increasing the slopes of the tuning curves and thus the discriminability of different stimuli.

To investigate whether the biases in the neural population also play a role in encoding prior information we divided the neurons into a group encoding more likely, or ‘expected’, stimuli and more unlikely, or ‘unexpected’ stimuli. Looking at the results from Fig 12, we selected the neurons with their maximal slope within one standard deviation of the prior probability distribution as those sensitive to ‘expected’ stimulus values and neurons with their maximal slope outside one standard deviation of the prior probability distribution as those sensitive to ‘unexpected’ stimulus values. The neurons sensitive to ‘expected’ stimuli had a stronger bias, or baseline activation, then neurons sensitive to ‘unexpected’ stimuli for all different prior distribution (Fig 14).

**Figure 14.**
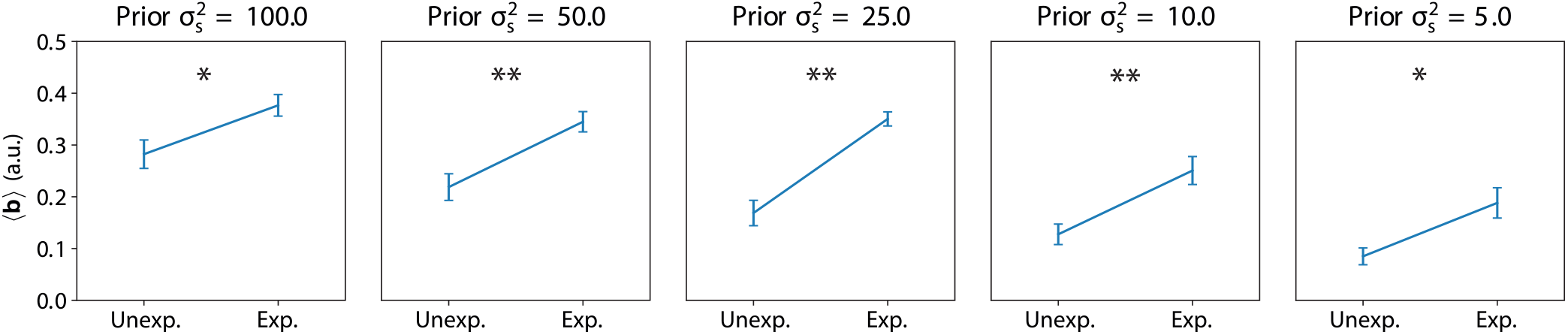
Bias influence by prior probability of stimuli. For every trained network we determined which neurons were sensitive to ‘expected’ stimuli and which were sensitive to ‘unexpected’ stimuli. The bias was much stronger for neurons sensitive to expected stimuli for all prior probability distributions. (* *p <* 0.02, ** *p <* 0.005).

## Discussion

### Summary of results

We have shown that a neural population is able to learn to incorporate prior knowledge optimally by error based learning without any probabilistic information, both for discrete priors (classification task) and continuous priors (stimulus estimation task). For both discrete and continuous priors, errors for likely stimuli are lower than errors for unlikely stimuli, as often observed in empirical literature. However, the mechanism through which this is achieved differs for both types of priors. In the classification task, a higher prior probability for a class led to higher activity in neurons encoding this class. On the other hand, stimuli with higher input contrast led to lower activity in the network. The increase in neural activity by the prior was caused by a lowered activation threshold of the neurons encoding the more likely class. When a contextual cue was added, a similar effect was achieved by a strong excitatory input to neurons encoding the more likely class and an inhibitory input to the neurons encoding the less likely class. In the stimulus estimation task the opposite effect of prior probability on the neural activity was observed. A narrower prior distribution led to lower activity in the neural population. The prior changed the distribution of tuning curves, increasing the density and narrowing the tuning curves. Not the peak of the tuning curves, but the maximal slope followed the prior distribution. Neurons with their maximal slopes around ‘expected’ stimuli values had a lower activation threshold then neurons with their maximal slope around ‘unexpected’ stimuli values. Thus the effects of prior probability on neural activations are affected by a range of different factors, like lower activation thresholds, higher tuning curve density or narrower tuning curve width, which can play a different role depending on the exact task that is being performed.

### Relation to previous work

These results give a strong indication that it is possible to learn an efficient incorporation of prior knowledge based on errors in individual observations, and without the need of higher cognitive regions keeping track of explicit probabilities. With this study we extend previous findings of^11^ to incorporate prior knowledge and show that even such unbalanced class observations lead to optimal Bayesian inference. We suggest a mechanism that uses changes in the activation thresholds to encode prior probabilities. Such changes of intrinsic neural properties have been proposed previously as an efficient mechanism to achieve sparse responses to stimuli^20^ and maximize information transmission. Interestingly, we find a similar mechanism to encode prior probabilities of stimuli optimally. Previous models describing the representations of uncertainty in the brain, have strongly focused on the role of spiking behaviour. The variability caused by such spiking behaviour could be connected to the mean firing rate of a neuron (in case of Poisson like neurons) where both encode the uncertainty of a stimulus^6^. Alternatively, the variability and the mean of neural responses can be representing separate aspects of a posterior probability distribution, with only the variability of neural firing representing uncertainty^7^. Our work shows that rate based models are also able to perform such probabilistic tasks requiring knowledge about prior distributions, shedding light on a completely different way of representing prior knowledge.

The learning through observations demonstrated by our network is supported by experimental studies showing the formation of an environmental prior over the lifetime^8,21^ and could potentially be how humans and animals learn to from optimal Bayesian priors^5, 22, 23^.

The representations of prior knowledge have been empirically studied in various settings. Conflicting observations reporting either increasing or decreasing neural responses to expected or unexpected stimuli have been found^14,24–30^. Here we showed that different factors can influence such neural responses, often indistinguishable from purely the average neural activity in a population. In the classification task, our model predicts stronger activity in response to expected stimuli, which has been observed in multiple studies^24,25,27^. However, in the estimation task, our model predicts lower activity in response to expected stimuli in line with experiments showing lower blood-oxygen-level-dependent (BOLD) responses in humans^14,28^ and electrophysiological responses in non-human primates^29,30^ to expected stimuli. Besides the particular task that is being performed, differences in neural activations in response to expected or unexpected stimuli can also result from the particular neurons that are measured during experiments. More detailed experimental measurements revealed that responses are higher in neurons selective to expected stimuli versus those not selective to expected stimuli^14,25^, similar to our results in Fig 4.

Another possible factor influencing the sparsity of responses could be the learning signal in response to unexpected stimuli. We have shown that while unexpected stimuli have sparser responses in the classification task, they lead to stronger learning signals, which could in turn lead to experimental measurements that show increased activity.

The properties of individual neurons, such as the activation threshold, gain, sharpness of the tuning curves or density of the tuning curves are all factors that determine the average neural activity of a population. We showed that all these factors can be influenced by the prior probability of a certain stimulus. Which factors are contributing strongest depends on the actual task that is being performed, leading to different effects on the neural activity.

Our model shows a clear role for the bias and to lesser extent the gain of individual neurons in the encoding of expected stimuli. Interestingly, lowered activation thresholds as suggested by our model have also been implied by observations that neurons encoding a stimulus fire already more strongly in response to a cue indicating that the stimulus is expected^31–33^, where the latter study also observed a subgroup of neurons showing gain effects in response to expected stimuli.

Besides the effects on the bias and gain of individual neurons we also found that the tuning curves of individual neurons adapted to efficiently encode more likely stimuli in the estimation task. The efficient allocation of limited neural resources to more likely stimuli has been suggested as a driving force underlying the formation of neural representations of prior knowledge^4,10,19,34,35^. Simulations have shown that maximizing Fisher information under limited resources results in denser and narrower tuning curves around expected stimuli values. Our results agree with such an efficient allocation of resources, with the important notion that the slopes of the tuning curves are actually concentrated around the most likely stimulus values. While some studies have suggested that the peak of a tuning curve is most important for encoding a stimulus^36^, others claim that the steepest slope of a tuning curve is most important^37,38^. Interestingly, a theoretical analysis of tuning curve information properties has indicated that maximal-slope encoding is most efficient for two-alternative-forced-choice tasks (such as our classification task) as well as an estimation task under reasonable levels of neuronal noise^39^, whereas peak-tuning-curve encoding has been shown to be more efficient only in an estimation task with high neuronal noise and a small population of neurons. The representation developed by our model agrees well with these findings and makes interesting suggestions for empirical validation.

### Predictions for empirical validation

We have shown that the particular effect that the prior has on the neural activity depends on the particular task that is being performed. Different mechanisms can give rise to the observed effects on neural activity. To verify which mechanisms are at play under which circumstances, more detailed experiments are necessary, that not only investigate the effects of prior knowledge on neural activations, but also properties of individual neurons such as their receptive fields and activation thresholds.

We find a large effect on the activation thresholds of neurons, with only a small effect due to gain modulations when there is a cue to indicate which stimuli are expected and which are unexpected. Changes in the activation threshold as a result of neural activity have been observed in real neurons ^18^, and could potentially play a role in the encoding of prior knowledge. This mechanism we observed also allows for flexibly adjusting this activation threshold by increasing the baseline input to a neuron. This enables the neural population to adjust its prior expectations based on the context. While such lowered activation thresholds have also been implicitly observed by stronger firing rates for neurons encoding a stimulus in response to a cue indicating that the stimulus is expected^31–33^, future studies could further confirm these changes in activation threshold. Validating this prediction requires intracellular recordings of sub-threshold activity, to see whether a contextual cue increases sub-threshold membrane potentials in neurons encoding stimuli that have a higher prior probability.

In the estimation task our model suggests that neural tuning curves are organized such that more neurons have their maximal slope around more likely stimuli. This could be validated in the by mapping tuning curves in response to oriented gratings in the visual cortex. An overrepresention of the cardinal axis due to prior exposure, as empirically observed^4^, suggests that more neurons should have the maximal slope of their tuning curves around the cardinal axis, while less neurons have their maximal slope around the oblique orientations.

### Limitations

Our approach builds on several approximations and assumptions that are important to clarify. Such assumptions can be potential limiting factors to the extension where the shown results hold.

We have chosen to model the input to our network as Poisson neurons, both out of practical considerations that for such neurons a Bayesian model can be analytically derived, but also for prevalent evidence showing (close to) Poisson like behaviour in the brain^40,41^. It is important to note that there also exists strong deviations from Poisson like behaviour, especially in some brain areas^42^. Also on small time scales such deviations occur^43^. Non Poisson like behaviour could allow for a decoupling between the mean and variance in the spike rate, allowing for an alternative encoding of uncertainty^7^. Such a code could also require a different mechanism of representing prior knowledge, which allows for interesting directions for future research.

To make it possible to compare our network to an optimal Bayesian model, we have resorted to relatively simple behavioral tasks. The binary classification task requires the learning of a linear optimal decision boundary, something that is relatively easy to learn. Whether a similar mechanism as found here plays a role in more complex tasks with natural stimuli requires further investigation in the future.

To train our network we used the backpropagation algorithm. There has been much debate regarding the biological plausibility of this learning algorithm, which has been criticized because of its biological implausibility^44^, for example regarding the non-locality of the information needed to update the weights, or the symmetry in the connectivity needed to backpropagate information in the network. Recent work has shown how, through relatively mild assumptions on e.g. dendritic segregating, patterns of spike timing, inhibitory microcircuits, short-term plasticity and feedback connections backpropagation can be achieved by biological networks^45^. For example, by allowing learning signals to travel through random feedback weights drops the requirement for symmetrical connectivity^46^ and by using segregated dendrites, it is prevented that confusion between learning signals and stimulus information occurs^47^. At the same time, models trained through backpropagation have been successful at explained actual brain responses and learning behaviour, indicating that the brain might use similar learning mechanisms to adapt synaptic connections^11,48–50^. It remains an open question whether the brain has found ways to implement some biological approximation of backpropagation, or whether it is a shear coincidence that deep learning models develop similar representations and mechanisms as the brain does. Further development on biological plausible implementations of the backpropagation algorithm is needed to resolve this question in the future. By validating predictions from work such as this we can further establish the explanatory power of mechanisms learned through the backpropagation algorithm, for the understanding of cortical activity.

### Future work

One important aspect of cognitive processing that is not included in our model, but that can also have a major effect on neural activations, is the role that feedback processing plays in the brain. A popular theory that describes how prior knowledge influences our perceptual processing is predictive coding. According to this theory, the brain forms predictions about incoming sensory information based on its internal prior knowledge. Since prior knowledge ‘explains away’ some of the sensory input through feedback, this also influences neural responses to expected stimuli class^14^. Since we used only feed-forward neural networks to model neural responses, this prevents feedback mechanisms from playing a role in encoding prior knowledge. An important step for future simulations would be to incorporate such feedback connections.

One way of doing this is through recurrent computations by adding a time component to our networks. Such a time component also allows for distinguishing neural activity in response to a cue, from neural activity in response to actual stimuli. This would allow to answer the question how a neural population could dynamically adjust to expected stimuli in order to improve sensitivity around expected stimuli values.

### Conclusions

We have shown that a model population of neurons can learn to correctly represent and incorporate prior knowledge, by only receiving feedback about the accuracy of their inference from trial-to-trial and without any probabilistic feedback. The prior probability of a stimulus affects the neural responses of the neural populations to this stimulus, where activations can either increase or decrease depending on the task. We show different mechanisms that influence the neural activations. We find that the activation threshold of neurons encoding a certain stimulus play an important role in representing its prior probability, with more likely stimuli having a lower activation threshold. In a task where estimating the exact stimulus value is important, more likely stimuli also lead to denser receptive field distributions and more narrow receptive fields, allocating computational resources such that information processing is enhanced for more likely stimuli. These results pose some clear and testable predictions about the neural representations of prior knowledge in commonly used experimental paradigms that can guide new experiments.

## Acknowledgements

This research was supported by VIDI grant number 639.072.513 of The Netherlands Organization for Scientific Research (NWO).

## Author contributions statement

S.Q. and M.G. conceived the experiments. S.Q. ran the simulations illustrated in the manuscript and prepared the figures. S.Q. and M.G. wrote the main manuscript text. S.B. and M.P. provided feedback for additional analysis. All authors reviewed the manuscript.

## Additional information

### Competing interests

The authors declare no competing interests.

## Notes

### Competing Interest Statement

The authors have declared no competing interest.

